## Appendix for "Digitalizing heterologous gene expression in Gram-negative bacteria with a portable on/off module"

by

Belén Calles, Angel Goñi-Moreno, and Víctor de Lorenzo

Table of Contents


| | |
|---|---|
| **Appendix Information** | Mathematical modelling of a digitalizer module merged with an inducible expression system |
| **Appendix Fig. S1.** | Individual effect of the strength of the repression of the different components of the digitalizer module |
| **Appendix Fig. S2.** | SBOL representation of the business cargo of plasmid pS238D•M. |
| **Appendix Fig. S3.** | Comparison of dose-responses of cells bearing the XylS/*Pm* expression device with or without or the digitalizer module |
| **Appendix Fig. S4.** | Deactivation kinetics of the digitalized and non-digitalized XylS/*Pm* device |
| **Appendix Fig. S5.** | Influence of the transcriptional repressor in the performance of the the digitalizer module |
| **Appendix Fig. S6.** | Kinetics of NIa protease activity upon expression though the XylS/*Pm* device carrying or not the digitalizer module |
| **Appendix Fig. S7.** | Kinetics of GFP expression from the AlkS/*PalkB* system |
| **Appendix Fig. S8.** | Influence of Hfq chaperone on the sRNA efficiency |
| **Appendix Fig. S9.** | Population heterogeneity of *P. putida* cells expressing GFP from digitalized or not versions of the AlkS/*PalkB* device |
| **Appendix Fig. S10.** | Performance of the digitalized XylS/*Pm* system expressing the colicin E3 toxin in *P. putida* |
| **Appendix Table S1.** | Strains and plasmids used in this study |
| **Appendix Table S2.** | Primers used in the PCR reactions |

### Mathematical modelling of a digitalizer module merged with an inducible expression system

The general organization of the genetically-encoded device under examination is sketched as follows:

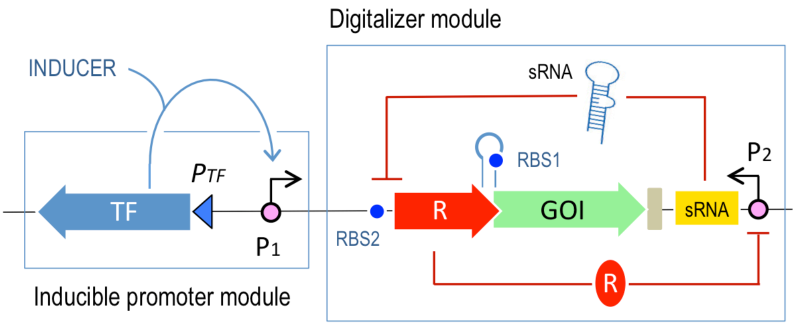


As explained in main body of the text, the parts of such a device include: [i] an inducible promoter $P_1$ with a given level of basal expression [ii] a strong, yet repressible promoter $P_2$ for transcription of a translation-inhibitory sRNA, [iii] a transcriptional repressor (R) expressed through the inducible promoter $P_1$ but translationally inhibited by the sRNA and [iv] a gene of interest (GOI) translationally coupled to the repressor gene. As shown in the scheme above, the repressor protein R targets the strong promoter $P_2$ from which the inhibitory sRNA is produced, so that R and sRNA are mutually inhibitory. The goal of this arrangement is to create a general-purpose genetic plug-in that minimizes the basal (i.e. non-induced) activity of $P_1$ and thus suppresses leaky gene expression levels. A key requirement of the design is that the device must be general-purpose e.g. the mechanism needs to be uncoupled from a specific $P_1$ promoter sequence, thus allowing users to plug-and-play their promoter of choice—which will be automatically digitalized by the device. This avoids hacking the operator regions of the promoter, an approach that would imply to re-design and re-engineer the function every time the user needed a different promoter.

As far as mathematical modelling is concerned, the above requirement means that the rates of binding and unbinding of a transcription factor (TF) to/from the promoter of choice cannot be altered. Nor can inducible transcription itself. The solution implies to add an external repression interaction to minimize basal levels, while making sure that such interaction vanishes after induction. In this way, the overimposed repression will not interfere with the intrinsic activity of the inducible promoter, thereby leading to maximization of the dynamic range at the output (i.e. difference between 0 and 1).

**Implementation of the circuit:** A explained in the main body of the article, the general circuit above was impemented in the shape of a plasmid construct with the following business DNA segment:

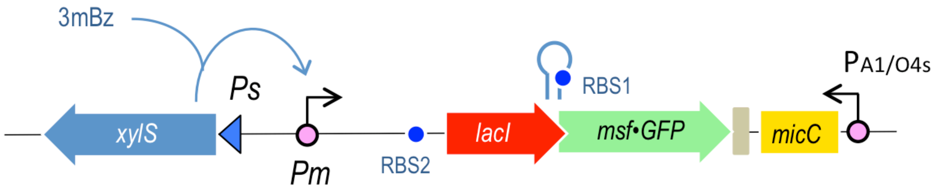


*Pm* is the promoter of interest (i.e., to digitalize), $XylS^a$ the transcription factor (in its active form) that induces *Pm* activity, $Pm^{XylSa}$ the promoter with the inducer bound, mRNA denotes the transcript that *Pm* triggers, LacI is the repressor translated from mRNA, GFP the fluorescent protein that is also translated from mRNA (both genes are transcriptionally coupled), $P_{A1/O4s}$ represents the repressible promoter, $P_{A1/O4s}{}^{LacI}$ denotes the repressible promoter with the repressor bound and sRNA is the translation inhibitory small RNA molecule. Inducer $XylS^a$ is constant through the simulations (i.e., $dXylS^a/dt=0$). Species are measured in molecules. Note that in the design shown above, external inhibition of the basal level comes from a sRNA molecule that is being express by a specific promoter ($P_{sRNA}$). Importantly, the expression of such sRNA was made conditional of the activity of the targeted promoter to be digitalised ($P_{target}$). This is, a regulator expressed by $P_{target}$ inhibits $P_{sRNA}$. Mathematical modelling was done to analyse the dynamical features of the system at different rate values. The performance of the switch was captured by the following set of ordinary differential equations:

$$\frac{dPm}{dt} = -k_3 \cdot XylS^a \cdot Pm + k_{-3} \cdot Pm^{XylS^a}$$

$$\frac{dPm^{XylS^a}}{dt} = k_3 \cdot XylS^a \cdot Pm - k_{-3} \cdot Pm^{XylS^a}$$

$$\frac{dmRNA}{dt} = k_4 \cdot Pm + k_5 \cdot Pm^{XylS^a} - k_6 \cdot mRNA - k_{10} \cdot sRNA \cdot mRNA$$

$$\frac{dLacI}{dt} = k_7 \cdot mRNA + k_{-11} \cdot PA1/O4s^{LacI} - k_{11} \cdot PA1/O4s \cdot LacI - k_{12} \cdot LacI$$

$$\frac{dPA1/O4s}{dt} = -k_{11} \cdot PA1/O4s \cdot LacI + k_{-11} \cdot PA1/O4s^{LacI}$$

$$\frac{dPA1/O4s^{LacI}}{dt} = k_{11} \cdot PA1/O4s \cdot LacI - k_{-11} \cdot PA1/O4s^{LacI}$$

$$\frac{dsRNA}{dt} = k_8 \cdot PA1/O4s - k_{10} \cdot sRNA \cdot mRNA - k_9 \cdot sRNA$$

$$\frac{dGFP}{dt} = k_7 \cdot mRNA - k_{13} \cdot GFP$$

Rates are described as follows: $k_1$ and $k_2$ are the generation (or entry) and degradation rates of $XylS^a$ molecules (not used in the model, since $XylS^a$ is constant), $k_3$ and $k_{-3}$ are the binding and unbinding, respectively, of $XylS^a$ to/from *Pm*, $k_4$ the basal transcription rate of *Pm*, $k_5$ is *Pm*'s induced transcription, $k_6$ is the degradation rate of mRNA, $k_7$ is the translation rate of LacI and GFP, $k_8$ the transcription rate of sRNA by $P_{A1/O4s}{}^{LacI}$, $k_9$ the degradation of sRNA, $k_{10}$ the repression from sRNA to mRNA, $k_{11}$ and $k_{-11}$ are the binding and unbinding, respectively, of LacI to/from $P_{A1/O4s}$, $k_{12}$ is the degradation rate of LacI, and $k_{13}$ is the degradation rate of GFP. We used standard rate values for modelling[1]. These are as follows: $k_3$ = 1 molecules$^{-1}$ hour$^{-1}$, $k_{-3}$ = 50 hour$^{-1}$, $k_4$ = 50 hour$^{-1}$, $k_5$ = 700 hour$^{-1}$, $k_6$ = 10 hour$^{-1}$, $k_8$ = 700 hour$^{-1}$, $k_9$ = 10 hour$^{-1}$, $k_{-11}$ = 70 hour$^{-1}$, and $k_{12}$ = 1 hour$^{-1}$. The next rates were modified within specific ranges to stress-test the model: $100 \leq k_7 \leq 120$ hour$^{-1}$, $0.2 \leq k_{10} \leq 14$ molecules$^{-1}$ hour$^{-1}$, $0.07 \leq k_{11} \leq 3.5$ molecules$^{-1}$ hour$^{-1}$, and $2.3 \leq k_{13} \leq 2.7$ hour$^{-1}$.

There are two key dynamical features at the core of the model that deserve specific attention. First, the DNA-binding repressor and the gene of interest (GOI) downstream of $P_{target}$ (*P1*) are both co-transcribed and translationally coupled. As a result, the sRNA inhibits both regulator and GOI at the same time. Secod, the *external* repression comes from an sRNA, instead of a protein, what reduces its action time. By avoiding translation the system works faster and reduces metabolic burden to the cell, which makes this device economically efficient.

[1] Miró-Bueno, J. M., & Rodríguez-Patón, A. (2011). A simple negative interaction in the positive transcriptional feedback of a single gene is sufficient to produce reliable oscillations. PloS One, 6(11), e27414.

**Simulation results.** The simulations shown in Fig. 3B and 3C of the main article body and Appendix Fig. S1 show that the double negative feedback loop (sRNA represses the regulator; the regulator inhibits the expression of sRNA) allows for digitalisation at specific parameter settings. The stronger that both regulations are, the more digital that $P_{target}$ response is. If any one of the regulations were weak, final performance would not show the required results. By digital we understand clear on/off states with a sharp transition in between. Moreover, the model suggested that, in the best scenario (i.e. strong repressions), only sRNA or mRNA will be found in the system at any given time, with the exception of a brief time lapse in which both molecules coexist before solving the issue into an stable state. Therefore, this returned a prediction: better to use a strong regulator (repressor) and strong sRNA. This was experimentally validated by using either LacI or LI as repressors (see main article body).

### Appendix Figures

**Appendix Figure S1. Individual effect of the strength of the repression of the different components of the digitalizer module**

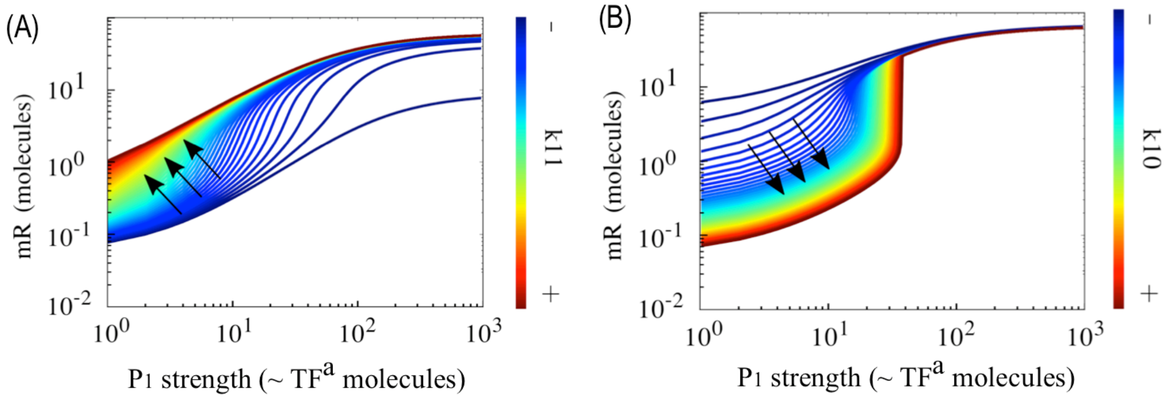


**(A)** Analysis of the transcriptional repressor component of the digitalizer module. The simulation considers different binding rates, which showed little effect in the system digitalization performance, although higher repressions correlated with more digital plots (indicated by arrows). **(B)** Simulations of the effect of the strength of the sRNA repression on the digitalization of the system showed a clear positive correlation. The stronger the repression the more digital the output of the device is. Arrows point to more digital-like performances. Although this component of the circuit seems to be the key, optimal digital behavior is achieved when combining strong repressions of both the sRNA and the repressor controlling its expression.

**Appendix Figure S2. SBOL representation of the business cargo of plasmid pS238D•M.**

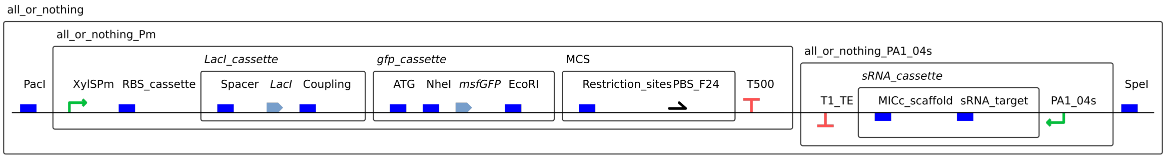


The design is focused on representing the hierarchical structure of the different components: One module contains *Pm* and its downstream parts, and another $P_{A1/04S}$ and its downstream parts. The complete SBOL design is available at SynBioHub [2] in https://synbiohub.org/user/Angel/Digitalizer/Digitalizer_collection/1/11824c40aa5636bf3c2da437d1f8a828ef7d64e3/share.

[2] McLaughlin *et al.* (2018). SynBioHub: A Standards-Enabled Design Repository for Synthetic Biology. *ACS Synthetic Biology*, *7*: 682-688.

**Appendix Figure S3. Comparison of dose-responses of cells bearing the XylS/*Pm* expression device with or without or the digitalizer module**

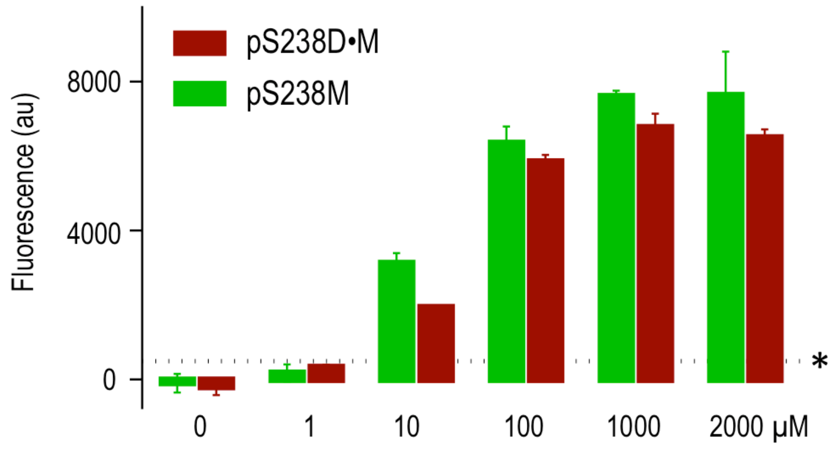


The dose-responses of *E. coli* cells harboring the XylS/*Pm* node either by itself (i.e. harboring the pS238M plasmid) or merged with the digitalizer module (i.e. harboring pS238D•M) were inspected as indicated in the legend of Fig. 1D of the main text. The GFP fluorescence signal was recorded in respect to the control strain (i.e. transformed with the empty pSEVA238 vector) for discarding auto-fluorescence. Cultures were grown in the absence and presence of increasing concentrations of inducer (3MBz) as indicated. Values below the horizontal dashed line lie within the non-confidence region of fluorescence detection in the instrument. Arbitrary units were normalized in all cases to the optical density of the cultures under examination (data for cells with pS238M are reused for comparison from the experiment of Fig. 1D of the main text).

**Appendix Figure S4. Deactivation kinetics of the digitalized and non-digitalized XylS/*Pm* device**

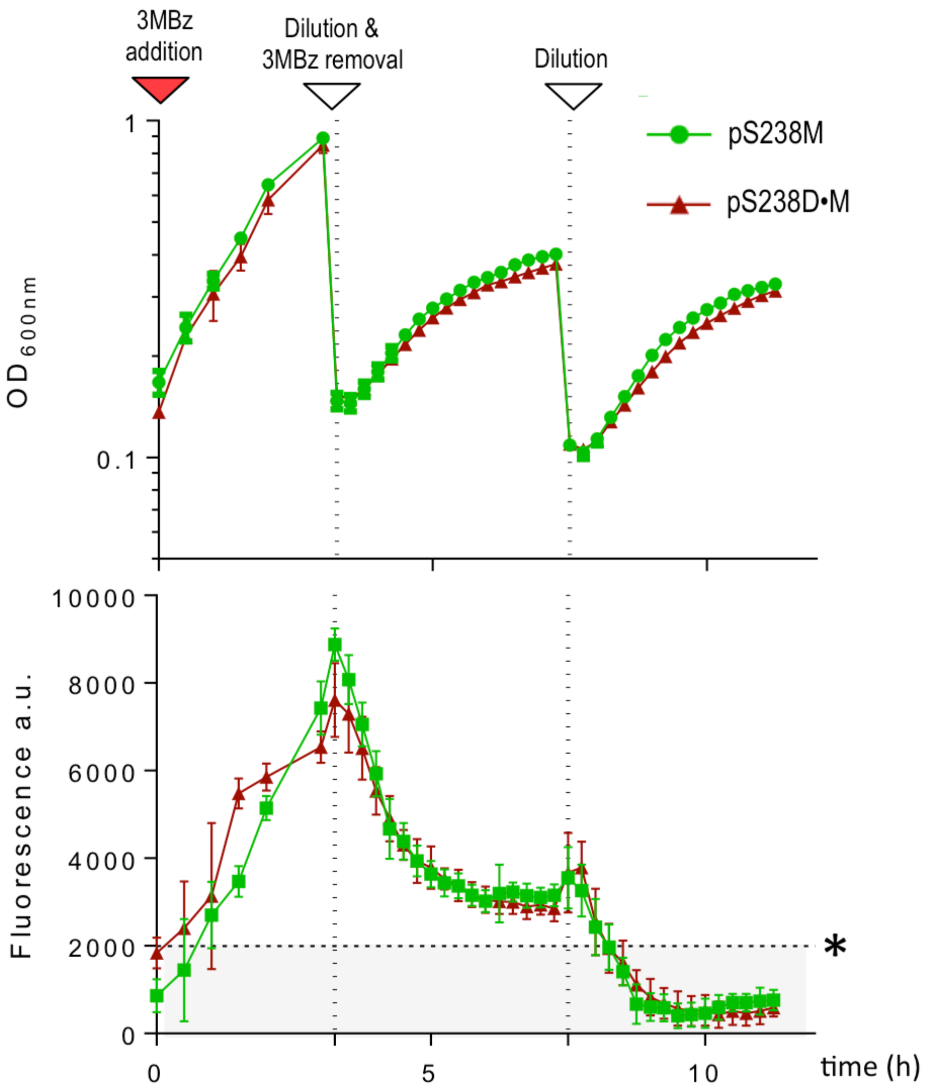


Growth curves (*upper* panel) and GFP fluorescence (lower panel) correspond to *E. coli* cells transformed either with pS238M (XylS/*Pm* node by itself, green) or with pS238D•M (digitalized version, (red plot). The experiment has 3 stages indicted by vertical dashed lines. As indicated, GFP induction at t = 0 was triggered by addition of 1 mM 3MBz, followed by two subsequent steps of 3MBz removal and re-dilution to mid-exponential growth phase ($OD_{600}$ ~ 0.4)/growth to stationary phase. Fluorescence was normalized to the corresponding $OD_{600}$s. Values below the horizontal dashed line lie within the non-confidence region of fluorescence detection in the instrument, as assessed with a control culture transformed with the empty plasmid pSEVA238.

**Appendix Figure S5. Influence of the transcriptional repressor in the performance of the the digitalizer module**

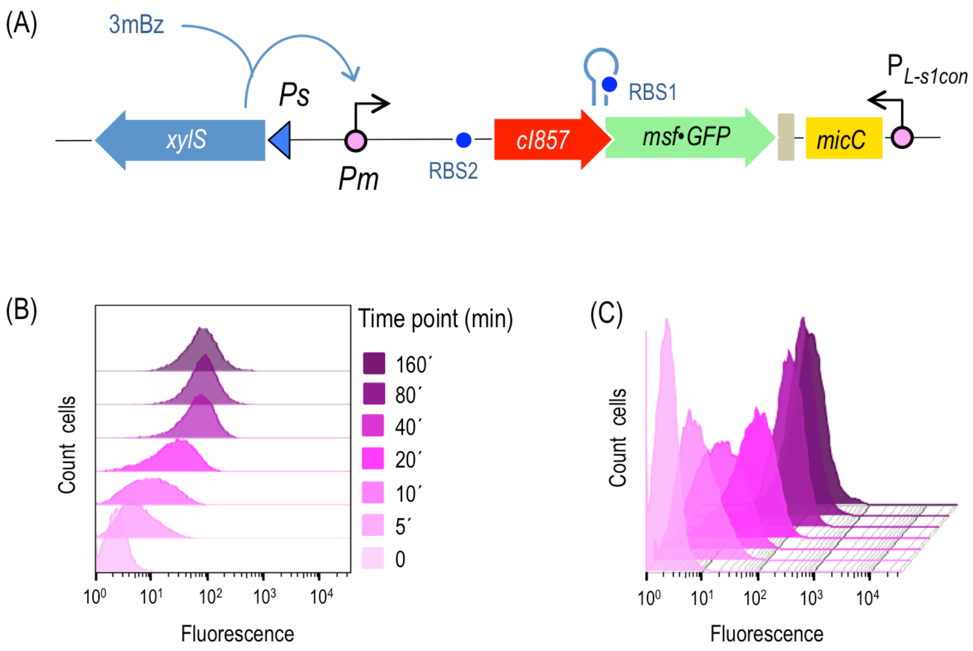


**(A)** Schematic representation of the digitalizer module harboring the thermosensitive variant of the CI repressor that is regulating a $P_{L\text{-}s1con}$ cognate promoter, driving the expression of a sRNA that was tailored to target, and inactivate the CI mRNA. **(B)** Kinetics of GFP expression in *E. coli* cells harboring the CI-dependent digitalizer module described above, analyzed by flow cytometry experiments in the absence (t=0) and the presence of a fix concentration of inducer (1.0 mM of 3MBz) at the indicated time points (5 to 160 min after induction).

**Appendix Figure S6. Kinetics of NIa protease activity upon expression though the XylS/*Pm* device carrying or not the digitalizer module**

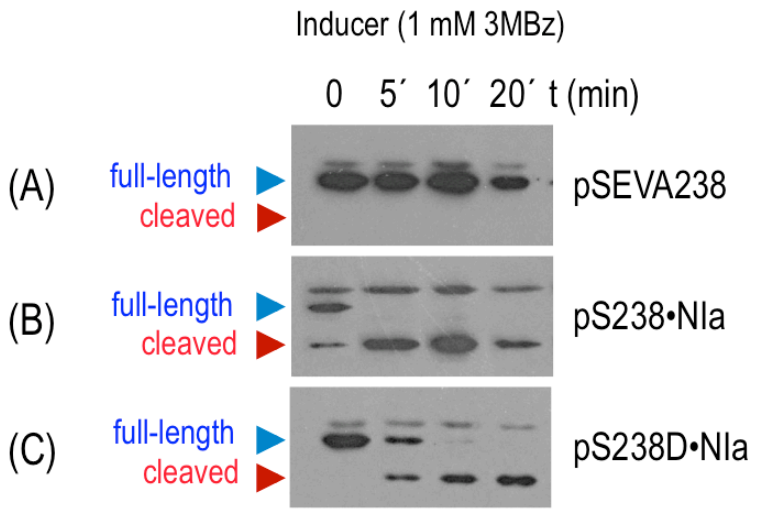


*E. coli* cells harboring a NIa sensitive and E-tagged variant of the TpiA protein (replacing the wild-type sequence) were transformed either with (A) pSEVA238, control with the XylS/*Pm* expression system, (B) pS238•NIa, expressing the NIa protease under the native XylS/*Pm* device and (C) pS238D•NIa the same but with the digitalized version of XylS/*Pm*. The cultures were grown until mid-exponential phase (t = 0) and NIa protease production was induced by addition of 1 mM 3MBz. Protein expresion was followed by Western blot assays of cell samples taken each 5 minutes after induction. As expected, the relative high basal level of pS238•NIa was detected as cleavage of at least half of the NIa-sensitive TpiA protein in the absence of inducer. In contrast, insertion of the digitalizer prevented leaky expression. As shown, NIa degraded TpiA protein between 5 and 10 min after induction. These results confirmed that the residual basal expression of XylS/*Pm* detectable only with this usper-sensitive method, was supressed when adding the digitalizing module to the expression device.

**Appendix Figure S7. Kinetics of GFP expression from the AlkS/*PalkB* system**

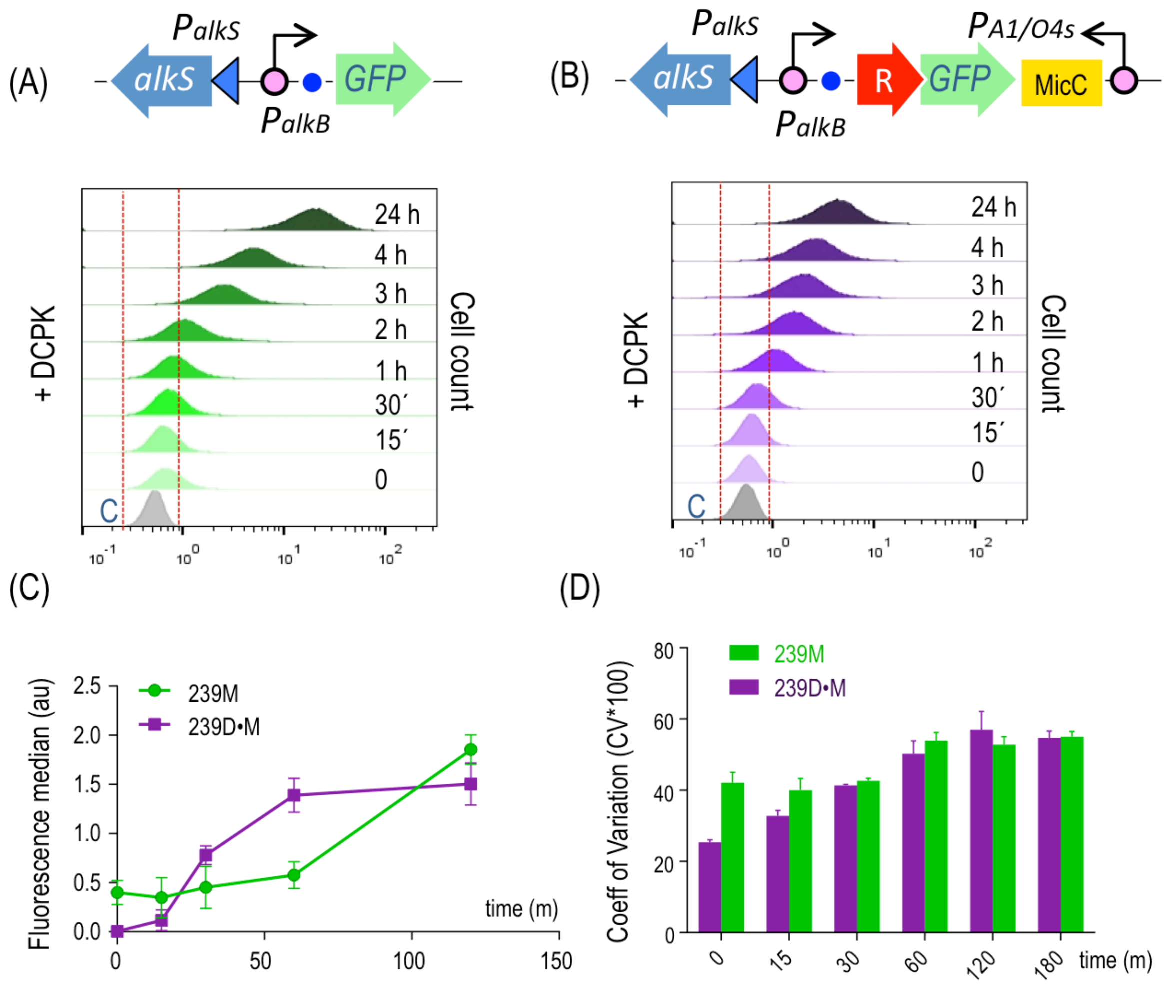


**(A)** Without digitalizer module, **(B)** With digitalizer module. Data were collected at the indicated time points, after induction with a fixed (1 mM) DCPK concentration. The host in both cases was *E. coli* CC118. Grey plot is the output of the strain containing a promotor-less GFP (pSEVA237M) and thus defines the region of no GFP fluorescence signal (comprised between red dashed lines). **(C)** Median expression of GFP from the AlkS/*PalkB* system (green plot) and its digitalized version (purple plot) before (t = 0) and a different time points after induction with DCPK, as indicated. Results include three independent experiments. **(D)** Population heterogeneity of cells analysed in the experiment previously described analysed by means of the coefficient of variation (percentage: CV*100).

**Appendix Figure S8. Influence of Hfq chaperone on the sRNA efficiency**

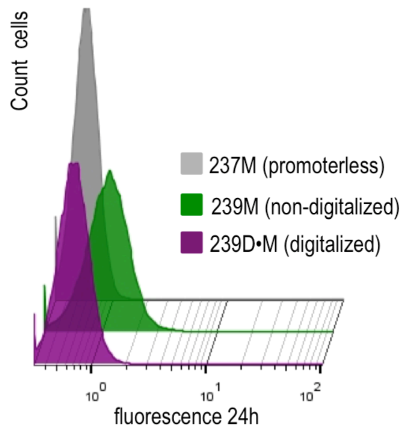


Comparison of the basal expression of GFP cloned in the AlkS/*PalkB* system (green plot) and its digitalized version (purple plot) analysed in an *E. coli* Hfq deficient strain. The lack of this RNA chaperone had little influence on the performance of the digitalizer module as the distribution of fluorescence in both non-fluorescence control strain (grey plot), containing a plasmid with the promotorless GFP, and cells harbouring the digitalized device (purple) virtually overlap.

**Appendix Figure S9. Population heterogeneity of *P. putida* cells expressing GFP from digitalized or not versions of the AlkS/*PalkB* device**

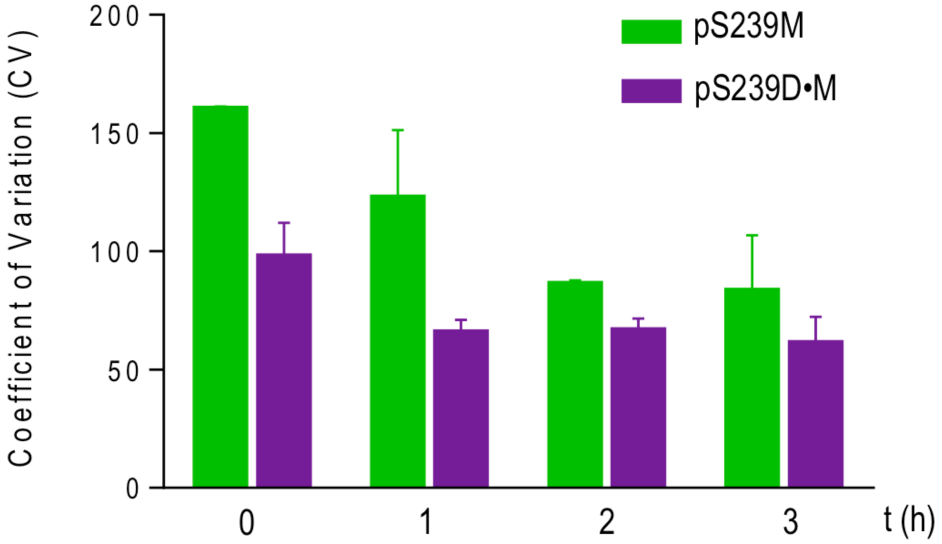


Coefficient of variation (CV) was determined in *P. putida* cells expressing GFP from digitalized (bearing pS239D•M, purple bars) or not (with pS239M, green bars) versions of the AlkS/*PalkB* device following induction with 1 mM DCPK as explained in Materials and Methods. Note the very significant increase in cell-to-cell homogeneity, especially at earlier induction times.

**Appendix Figure S10. Performance of the digitalized XylS/*Pm* system expressing the colicin E3 toxin in *P. putida***

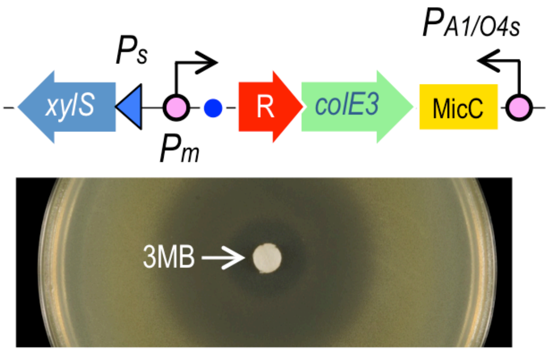


The image shows culture of a KT2440 *P. putida* strain transformed with plasmid pS438D•colE3 harbouring the highly toxic *colE3* gene under the control of the XylS/*Pm* digitalized device (skteched on top) and laid with soft agar on the surface of LB agar. The inducer of the system (0.5 M 3MBz) was added in a 3MM paper in the middle of the plate. A considerable halo of growth inhibition appeared as consequence of the action of the toxin in cells surrounding the inducer. On the contrary, cells far enough were able to grow, indicating that the basal expression is negligible in hosts other than *E. coli* as well.

**Appendix Table S1.** Strains and plasmids used in this study

| Strain or plasmid | Description | References and/or source |
|---|---|---|
| *E. coli* strains | | |
| JW4130 | Δ(*araD-araB*)567, Δ*lacZ4787*(*::rrnB-3*), $\lambda^-$, *rph-1*, Δ(*rhaD-rhaB*)568*, Δhfq-722::kan*, *hsdR514* | CGSC, Keio Collection (Baba et al, 2006) |
| MDS42 | Reduced-genome IS- free *E. coli* K12 strain with 704 genes deleted with respect to MG1655 parental strain | (Pósfai et al, 2006) |
| CC118 | $F^-$, Δ(ara-leu)7697, *araD139, Δ(lac)X74, phoAΔ20, galE, galK, thi, rpsE, rpoB, argE(Am), recA1* | (Manoil & Beckwith, 1985) |
| CC118 immE3 | CC118 strain with a chromosomal insertion of a mini-Tn*5* transposon constitutively expressing ImmE3 immunity protein, $Km^R$ | (Diaz et al, 1994) |
| W3110 Δ*tpiA* | Prototrophic derivative of wt K12 W3110 strain, $F^-$, $I^-$, IN *(rrnD-rrnE)1, rph-1, ΔtpiA* | Calles, *et al.* In preparation |
| *P. putida* strains | | |
| KT2440 | Prototrophic wild-type *P. putida* strain | (Nelson et al, 2002) |
| EM42 | KT2440 derivative; Δprophage1, Δprophage4, Δprophage3, Δprophage2, ΔTn7, Δ*endA-1,* Δ*endA-2* Δ*hsdRMS*, Δflagellum, ΔTn*4652* | (Martínez-García et al, 2014) |
| Plasmids | | |
| pSEVA226 | Derivative of pSEVA221 bearing a promoterless *luxCDABE* operon, $Km^R$; ori RK2, standard multiple cloning site | (Silva-Rocha et al, 2012) |
| pSEVA226-XylS/Pm | Derivative of pSEVA226 harboring the XylS/*Pm* expression system | (Silva-Rocha et al, 2012) |
| pSEVA231 | $Km^R$; *oriV* pBBR1, standard multiple cloning site | (Silva-Rocha et al, 2012) |
| pSEVA238 | Inducible expression vector; $Km^R$, *oriV* (pBBR1); xylS/*Pm*; standard multiple cloning site | This work |

| | | |
|---|---|---|
| pSEVA237M | Km$^R$; oriV(pBBR1), promoterless *msf•GFP* | (Benedetti et al, 2016) |
| pS238M | pSEVA238 derivative bearing the *msf•GFP* gene; Km$^R$; oriV (pBBR1); cargo (*xylS-Pm → msf•GFP*) | This work |
| pGA-LacI | Devirative of plasmid p57 inserted with a synthetic fragment containing *msf•GFP* gene and sRNA expression module. Ap$^R$; oriPMB1; *lacZ* | GeneArt® |
| pS238D•M | Derivative of pSEVA238 harboring harboring *msf•GFP* gene under the control of the LacI-dependent digitalizing module | This work |
| pPPVS20 | pSU8 derivative containing the the 3 ' terminal region of PPV cDNA from nt 3627 with NIa protease activity. P15A ori; Cm$^R$ | (Garcia et al, 1989) |
| pS238•NIa | pSEVA238 derivative bearing the *nIa* gene; Km$^R$; oriV (PBBR1); cargo (*xylS/Pm → nIa*) | This work |
| pBCL3-57-NIa | pUC18 derivative bearing a C-terminal E-tagged *tpiA* gene inserted with the NIa protease recognition sequence at residue E57 | Calles *et al.* in preparation |
| pS238D•NIa | Derivative of pSEVA238 harboring the potyvirus *nIa* protease gene under the control of the LacI-dependent digitalizing module | This work |
| pSEVA438 | Inducible expression vector; Sm$^R$; *oriV* (pBBR1); xylS/*Pm*; standard multiple cloning site | This work |
| pEDF5 | pVLT derivative expressing *colE3* gene. Tc$^R$; *oriV* RSF1010 | (Diaz et al, 1994) |
| pS438D•colE3 | Derivative of pSEVA438 harboring the *colE3* gene harboring *msf•GFP* gene under the control of the LacI-dependent digitalizing module | This work |
| pSEVA429 | Inducible expression vector; Sm$^R$; *oriV* (RK2); AlkS/$P_{AlkB}$; standard multiple cloning site | (Martínez-García et al, 2015) |
| pS239M | pSEVA239 derivative bearing the *msf•GFP* gene; Km$^R$; oriV (PBBR1); cargo (AlkS/$P_{AlkB}$ → *msf•GFP* ) | This work |
| pS239D•M | Derivative of pSEVA239 harboring harboring the AlkS/$P_{AlkB}$ gene under the control of the LacI-dependent digitalizing module | This work |

| | | |
|---|---|---|
| pGA-CI | Derivative of plasmid pMK-RQ containing a synthetic DNA fragment with $cI_{857}$ gene and sRNA expression module. $Km^R$; *oriV* Col E1 | GeneArt® |
| pS238D1•M | Derivative of pSEVA238 harboring the *msf•GFP* gene under the control of the CI-dependent digitalizing module | This work |

**Appendix Table S2.** Primers used in the PCR reactions a)

| Oligonucleotide | Sequence |
|---|---|
| LacI-F | CAACAGAAACAATAATAATGGAGTCATGACCATGCCTAGGAGGAAAAAACATATGGTGAAACCAGTAACGTTATAC |
| LacI-R2 | CATTATCATCACCACCATCCTAATGATGGTGGTGATGATGCTGCCCGCTTTCCAGTCGGG |
| GFP -F2 | CAGCATCATCACCACCATCATTAGGATGGTGGTGATGATAATG |
| GFP-R | GTTCTGAGGTCATTACTGGATCTATCAACAGGAGTCCAAGACTAGTTTTATCAAAAAGAGTG |
| msfGFPI-F | ACGTCCAAGCTTAGGAGGAAAAAACATATGGCTAGCCGTAAAGGTGAAG |
| msfGFP-R | GATATACTAGtTTATTTGTAGAGTTCATCCATG |
| JB653-F | GCGATGAGCTCAAAGAGGAGAAATTAAGCATGAGTAAATCACTGTTTAGAGGCC |
| JB653-R | CGTCGGGTACCTTACTGAGTGTAAACAAATTCCCCATC |
| NIa-F SR syst | CGTCGGCTAGCAGTAAATCACTGTTTAGAGGCCTG |
| ColE3-F | ACGTACGCTAGCGGTGGCGATGGACGCGGCCATAAC |
| ColE3-R | ACGTACGAATTCTCAAAGATATTTCTTGATATTTCG |

a) Sequences of oligonucleotides employed in this study. Restriction sites entered for cloning purposes are underlined.

REFERENCES


Baba T, Ara T, Hasegawa M, Takai Y, Okumura Y, Baba M, Datsenko KA, Tomita M, Wanner BL, Mori H (2006) Construction of Escherichia coli</em> K-12 in-frame, single-gene knockout mutants: the Keio collection. *Mol Sys Biol* **2:** 2006.0008

Benedetti I, Nikel PI, de Lorenzo V (2016) Data on the standardization of a cyclohexanone-responsive expression system for Gram-negative bacteria. *Data in Brief* **6:** 738-744

Diaz E, Munthali M, Lorenzo V, Timmis KN (1994) Universal barrier to lateral spread of specific genes among microorganisms. *Mol Microbiol* **13:** 855-861

Garcia JA, Riechmann JL, Martín MT, Laín S (1989) Proteolytic activity of the plum pox potyvirus NIa-protein on excess of natural and artificial substrates in *Escherichia coli. FEBS Lett* **257:** 269-273

Manoil C, Beckwith J (1985) TnphoA: a transposon probe for protein export signals. *Proc Nat Acad Sci USA* **82:** 8129-8133

Martínez-García E, Aparicio T, Goñi-Moreno A, Fraile S, de Lorenzo V (2015) SEVA 2.0: an update of the Standard European Vector Architecture for de-/re-construction of bacterial functionalities. *Nucl Acids Res* **43:** D1183-D1189

Martínez-García E, Nikel PI, Aparicio T, de Lorenzo V. (2014) *Pseudomonas* 2.0: genetic upgrading of P. putida KT2440 as an enhanced host for heterologous gene expression. *Microb Cell Fact* 13: 159.

Nelson KE, Weinel C, Paulsen IT, Dodson RJ, Hilbert H, Martins dos Santos VAP, Fouts DE, Gill SR, Pop M, Holmes M, Brinkac L, Beanan M, DeBoy RT, Daugherty S, Kolonay J, Madupu R, Nelson W, White O, Peterson J, Khouri H et al (2002) Complete genome sequence and comparative analysis of the metabolically versatile *Pseudomonas putida* KT2440. *Environ Microbiol* **4:** 799-808

Pósfai G, Plunkett G, Fehér T, Frisch D, Keil GM, Umenhoffer K, Kolisnychenko V, Stahl B, Sharma SS, de Arruda M, Burland V, Harcum SW, Blattner FR (2006) Emergent Properties of Reduced-Genome *Escherichia coli. Science* **312:** 1044-1046

Silva-Rocha R, Martínez-García E, Calles B, Chavarría M, Arce-Rodríguez A, de las Heras A, Páez-Espino AD, Durante-Rodríguez G, Kim J, Nikel PI, Platero R, de Lorenzo V (2012) The Standard European Vector Architecture (SEVA): a coherent platform for the analysis and deployment of complex prokaryotic phenotypes. *Nucl Acids Res* **41:** D666-D675
